# Deep neural language modeling enables functional protein generation across families

**DOI:** 10.1101/2021.07.18.452833

**Authors:** Ali Madani, Ben Krause, Eric R. Greene, Subu Subramanian, Benjamin P. Mohr, James M. Holton, Jose Luis Olmos, Caiming Xiong, Zachary Z. Sun, Richard Socher, James S. Fraser, Nikhil Naik

**Author notes:** Correspondence to: Ali Madani, Nikhil Naik.

## Abstract

Bypassing nature’s evolutionary trajectory, *de novo* protein generation—defined as creating artificial protein sequences from scratch—could enable breakthrough solutions for biomedical and environmental challenges. Viewing amino acid sequences as a language, we demonstrate that a deep learning-based language model can generate functional artificial protein sequences across families, akin to generating grammatically and semantically correct natural language sentences on diverse topics. Our protein language model is trained by simply learning to predict the next amino acid for over 280 million protein sequences from thousands of protein families, without biophysical or coevolutionary modeling. We experimentally evaluate model-generated artificial proteins on five distinct antibacterial lysozyme families. Artificial proteins show similar activities and catalytic efficiencies as representative natural lysozymes, including hen egg white lysozyme, while reaching as low as 44% identity to any known naturally-evolved protein. The X-ray crystal structure of an enzymatically active artificial protein recapitulates the conserved fold and positioning of active site residues found in natural proteins. We demonstrate our language model’s ability to be adapted to different protein families by accurately predicting the functionality of artificial chorismate mutase and malate dehydrogenase proteins. These results indicate that neural language models successfully perform *de novo* protein generation across protein families and may prove to be a tool to shortcut evolution.

---

Protein engineering can have a transformative impact on biomedicine, environmental science, and nanotechnology, among other fields. Traditional evolutionary protein engineering methods perform iterative screening/selection to identify proteins with desired functional and structural properties. In contrast, rational or *de novo* protein design methods aim to create artificial proteins from scratch, with the goal of improved efficiency and precision. Structure-based *de novo* design methods^1–5^ employ biophysical principles to design proteins that will fold into a desired structure using simulation-based methods and statistical analysis of experimental data. Coevolutionary methods^6–9^ build statistical models from evolutionary sequence data to specify novel sequences with desired function or stability. Both structural and coevolutionary approaches are not without limitations. Structural methods rely on scarce experimental structure data and difficult or intractable biophysical simulations^3,10^. Coevolutionary statistical models are tailored to specific applications, frequently rely on multiple sequence alignments, and struggle to operate in underexplored regions of protein landscape^10^. Recently, deep neural networks have shown promise as generative and discriminative models for protein science and engineering^11–15^. Their ability to learn complex representations could be essential to effectively utilize an exponentially growing source of diverse and relatively unannotated protein data—public databases containing millions of raw unaligned protein sequences^16–18^, enabled by the dramatic reduction in sequencing costs.

Inspired by the success of deep learning-based natural language models (LMs) trained on large text corpora that generate realistic text with varied topics and sentiments^19–23^, we develop ProGen, a protein language model trained on millions of raw protein sequences that generates *de novo* artificial proteins across multiple families and functions. While prior work has shown that natural language-inspired statistical representations of proteins are useful for protein informatics tasks such as stability prediction, remote homology detection, and secondary structure prediction^10,24–26^, we show that the latest advances in deep learning-based language modeling can be adopted to generate artificial protein sequences, from scratch, that function as well as natural proteins.

ProGen is iteratively optimized by learning to predict the probability of the next amino acid given the past amino acids in a raw sequence, with no explicit structural information or coevolutionary assumptions. Trained in this unsupervised manner from a large, varied protein sequence database, ProGen learns a universal, domain-independent representation of proteins that subsumes local and global structure motifs and higher-order residue interactions, analogous to natural language models learning semantic and grammatical rules. After training, ProGen can be prompted to generate full-length protein sequences for any protein family from scratch, with a varying degree of similarity to natural proteins.

ProGen is a 1.2 billion-parameter neural network trained using a publicly available dataset of 280 million protein sequences. A key component of ProGen is conditional generation^23,27–29^: sequence generation controlled by property tags provided as input to the language model. In the case of natural language, these control tags may be style, topics, dates, and other entities (Fig. 1a). For proteins, the control tags are properties such as protein family, biological process, and molecular function, that are available for a large fraction of sequences in public protein databases (Fig. 1b, Fig. S1). By specifying one or more control tags (e.g., Protein Family: Pfam ID PF16754, Pesticin) and none or a few amino acids at the beginning of a sequence, we can significantly constrain the sequence space for generation and improve generation quality, as shown by our prior computational experiments^30^.

**Figure 1:**
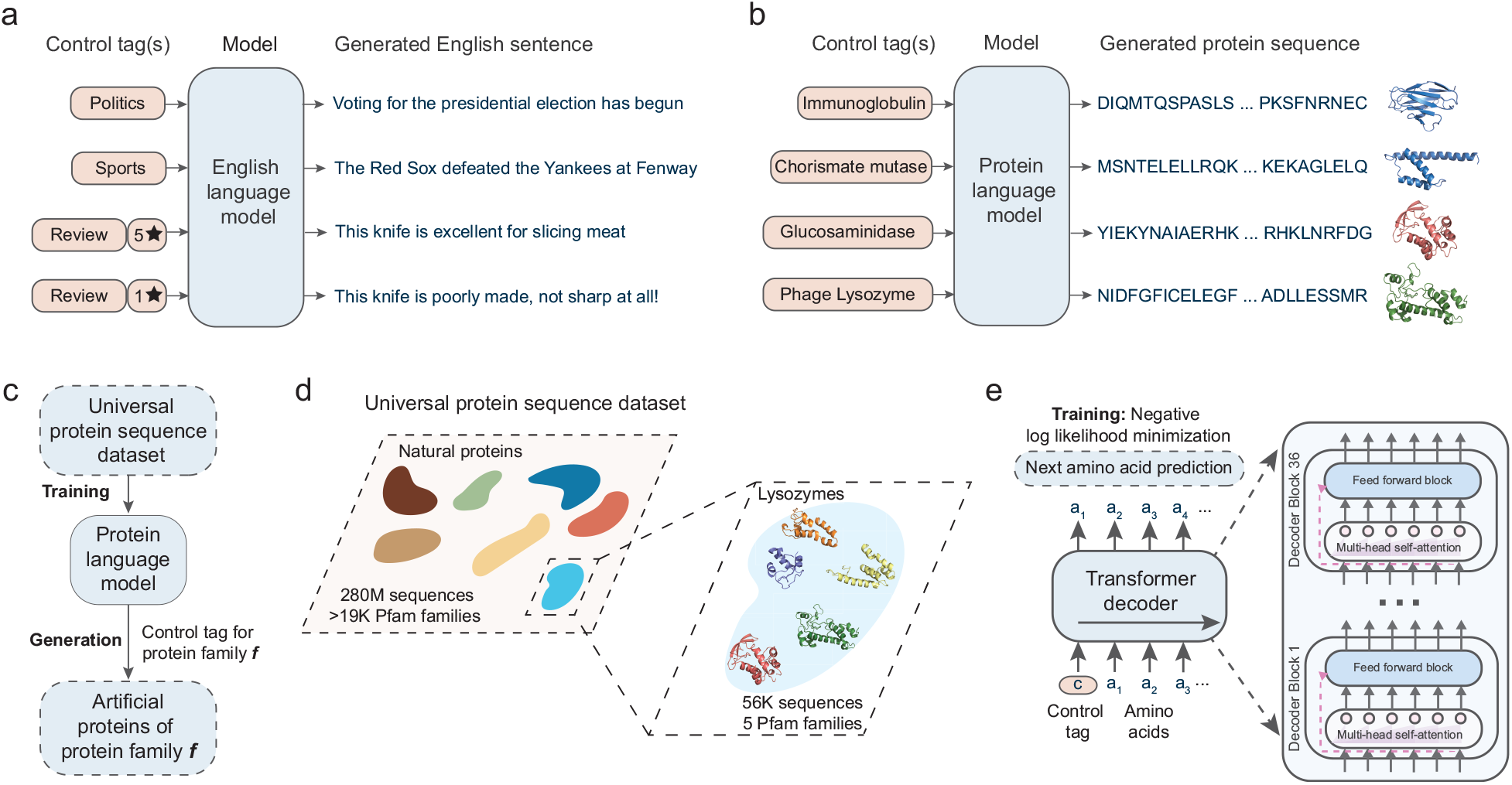
De novo protein generation with conditional language modeling. (a) Conditional language models are deep neural networks that can generate semantically and grammatically correct, yet novel and diverse natural language text, steerable using input control tags that govern style, topic, and other entities. Analogous to natural language models, we develop ProGen, a conditional protein language model (b) that generates diverse artificial protein sequences across protein families based on input control tags (c). ProGen is trained using a large, universal protein sequence dataset (d) of 280 million naturally-evolved proteins from thousands of families, of which five diverse lysozyme families are experimentally characterized in this study. ProGen is a 1.2 billion parameter neural network (e) based on the Transformer architecture which utilizes a self-attention mechanism for modeling comprehensive residue-residue interactions. ProGen is trained to generate artificial sequences by minimizing the loss over the next amino acid prediction problem on the universal protein sequence dataset.

Let *a* = (*a*_1_, …, *a*_*na*_) be a sequence of amino acids that specifies a protein of length *n*_*a*_ − 1 appended with an “end of sequence” token. Let *c* = (*c*_1_, …, c_*nc*_) be an associated set of descriptors such as protein family or source organism, i.e., ‘control tags’, through which we would like to control generation of amino acid sequences. Let *x* = [*c*; *a*] be the sequence formed by prepending a control tag sequence to an amino acid sequence. The probability over such a combined sequence of length *n* = *n*_*a*_ + *n*_*c*_ is then *p*(*x*). Language modeling decomposes the problem of generating *x* into a next-token prediction problem^31^, where a token can either be an amino acid or a control tag. We train a neural network with parameters *θ* to minimize the negative log-likelihood over a dataset *D* = {*x*^1^, …, *x*^|*D*|^}

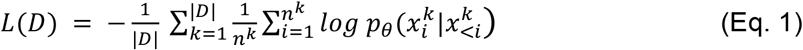

A new protein *<u>a</u>* of length *m*_*a*_ with desired properties encoded by a control tag sequence *<u>c</u>* of length *m*_*c*_ can then be generated by sequentially sampling its constituent tokens:*p*_*θ*_ (*a*_1_ | <u>c</u>), *p*_*θ*_ (*a*_2_ | <u>*a*_1_</u>, <u>*c*</u>), …, *p*_*θ*_ (*a*_*j*_ | *a*_<*j*_, <u>*c*</u>) ^30^. Generation continues until the model generates an “end of sequence” token.

We utilize a state-of-the-art transformer-based^19^ neural network architecture for constructing ProGen. The transformer learns long-range context within sequences using a series of stacked layers, each containing a self-attention mechanism (Fig. 1e). The self-attention mechanism in each layer infers pairwise interaction relationships between all positions in its input sequence^32^. Stacking multiple self-attention layers allows us to go beyond lower-order residue correlations to learn complex higher-order interactions. In contrast to transformer-based language models that encode amino acid sequences for discriminative protein prediction tasks^25,33,34^, ProGen is a decoder transformer tailored for autoregressive generation.

ProGen’s transformer architecture has 36 layers, and 8 self-attention heads per layer and a total of 1.2 billion trainable neural network parameters (see Methods). To train ProGen, we collected a universal protein sequence dataset containing 280 million protein sequences (from >19,000 Pfam^18^ families) and associated metadata (as control tags) from UniParc^16^, UniprotKB^17^, Pfam^18^, and NCBI taxonomic information^35^ (Fig. 1d, Table S1). We trained ProGen to minimize the negative log-likelihood defined in Eq. 1 using this dataset with a batch size of 2048 for one million iterations. Training was performed across 256 Google Cloud TPU v3 cores for two weeks. Once trained, ProGen could be used to generate novel protein sequences from scratch by specifying a control tag (e.g. protein family identifier from Pfam), Fig. 1c.

We experimentally evaluated ProGen’s ability to generate functional artificial amino acid sequences by testing its generations from the lysozyme clan^18,36^. Lysozymes are antibacterial enzymes that disrupt the integrity of the cell wall by catalyzing the hydrolysis of 1,4-beta-linkages between N-acetylmuramic acid and N-acetyl-D-glucosamine residues. Within the lysozyme clan, we chose to work with five families that lack disulfide bonds as to be amenable to high-throughput cell-free expression: Phage lysozyme, Pesticin, Glucosaminidase, Glycoside hydrolase, and Transglycosylase. These lysozyme families contain significant protein sequence diversity (Table S3) with average sequence length varying between 84-167 across families. The sequences also show large structural diversity and multiple structural folds (Fig. S2), which represents a challenging design space for a model trained without any explicit structural information. We collected a dataset of 55,948 sequences from these five families from Pfam and UniprotKB sources for obtaining positive controls and for fine-tuning^37^ ProGen. Fine-tuning involves making limited, computationally inexpensive, gradient updates to the parameters of the trained model (see Methods) to further improve its ability to generate protein sequences in the local lysozyme sequence neighborhood^38^.

After fine-tuning ProGen using the curated lysozyme dataset, we generated one million artificial sequences using ProGen by providing the Pfam ID for each family as a control tag. Our artificial lysozymes span the sequence landscape of natural lysozymes, Fig. 2a, across five families which contain diverse protein folds, active site architectures, and enzymatic mechanisms^39,40^. As our model can generate full-length artificial sequences within milliseconds, a large database can be created to expand the plausible sequence diversity beyond natural libraries (Table S3). Although artificial sequences may diverge from natural sequences purely from a sequence identity calculation, (Fig. 2b, Fig. S3), they demonstrate similar residue position entropies when forming separate multiple sequence alignments of natural and artificial proteins within each family, Fig. 2c. This indicates that the model has captured evolutionary conservation patterns without training on explicit alignment information such as with Potts models^41^, as implemented in direct coupling analysis^6,42^.

**Figure 2:**
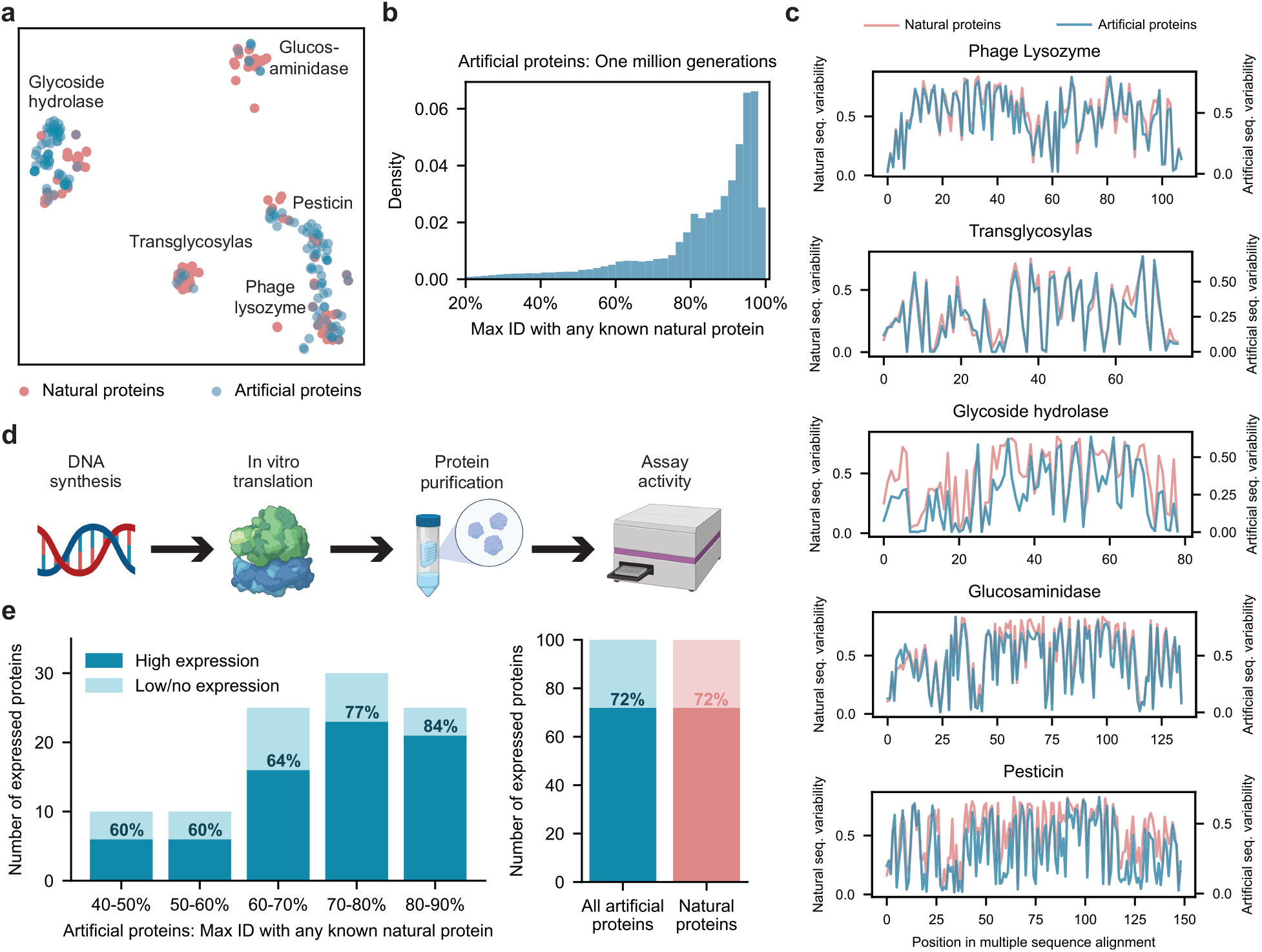
Generated artificial antibacterial proteins are diverse and express well in our experimental system. When analyzed using t-SNE, artificial sequences from our model are shown to span the landscape of natural proteins from five lysozyme families (a). Each point represents a natural or generated sequence embedded in a two-dimensional t-SNE space. With sufficient sampling, ProGen can generate sequences that are highly dissimilar from natural proteins (b). Max ID measures the maximum identity of an artificial protein with any publicly available natural protein. (c) Artificial proteins maintain similar evolutionary conservation patterns as natural proteins across families. Plots demonstrate the variability at each aligned position for a library of proteins. Conserved positions are represented as curve dips. From our generated proteins, we select one hundred proteins for synthesis and characterization in our experimental setup (d). Artificial proteins express well even with increasing dissimilarity from nature (40-50% max ID) and yield comparable expression quality to one hundred representative natural proteins (e).

To experimentally evaluate ProGen performance across a range of sequence divergences from natural proteins, we selected one hundred sequences filtered based on generation quality and diversity to natural sequences, measured as top-hit identities (“max ID”) to any protein in our training dataset containing 280 million proteins, which is primarily composed of UniParc^16^ (Fig. S4). We first ranked sequences for generation quality using a combination of an adversarial discriminator^22,43^ and model log-likelihood scoring^44^ (see Methods) and then sampled from top-ranked sequences to span a range of max IDs between 40-90%. We also ensured that no two generated samples share over 80% identity with each other. Our final selection included 100 artificial sequences (Table S2), with a minimum of 8 proteins from each protein family. The average sequence length for artificial proteins varies between 93-179 across families, comparable to natural lysozymes in our curated dataset from Pfam. We also selected a positive control group from the 55,948 curated lysozyme sequences. We clustered the natural sequences with mmseqs2^45^ and chose roughly 20 cluster-representative sequences from each of the five families (see Methods).

To evaluate function, full-length genes were synthesized and purified via cell-free protein synthesis and affinity chromatography (see Methods). In the positive control set of 100 natural proteins, 72% were well-expressed based on chromatography peaks and band visualization. The ProGen-generated proteins express equally well (72/100 total) across all bins of sequence identity to any known natural protein (maxID 40-90%), Fig 2e. In addition, we designed artificial proteins using bmDCA^6^, a statistical model based on direct coupling analysis, which explicitly approximates first and second-order residue dependencies. Although trained using publicly available code on the same sequences as ProGen and additional alignment information (see Methods), bmDCA was unable to fit three out of the five lysozyme families, and exhibited 60% detectable expression (30/50 proteins) for the remaining two protein families. These results indicate that ProGen can generate artificial proteins that are structurally well-folded for proper expression as compared to a batch of natural proteins, even when sequence alignment size and quality limit the success of alternative approaches.

Next we examined activity based on quench release of fluorescein labeled *Micrococcus lysodeikticus* cell wall (Molecular Probes EnzChek Lysozyme kit) using 90 randomly chosen proteins out of each expressed set of 100. Proteins were prepared in 96-well plate format to extract fluorescence curves over time, Fig 3a. Hen egg white lysozyme (HEWL), a naturally-evolved exemplar protein, was measured as positive control, in addition to ubiquitin as negative control. Proteins that generated fluorescence one standard deviation above the maximum fluorescence of any negative control were considered functional. Among our artificial proteins, 73% (66/90) were functional and exhibited high levels of functionality across families, Fig. 3c. The representative natural proteins exhibited similar levels of functionality with 59% (53/90) of total proteins considered functional. None of the bmDCA artificial proteins exhibited a detectable level of functionality. These results indicate that ProGen generates protein sequences that not only can express well but also maintain enzymatic function for diverse sequence landscapes across protein families.

**Figure 3:**
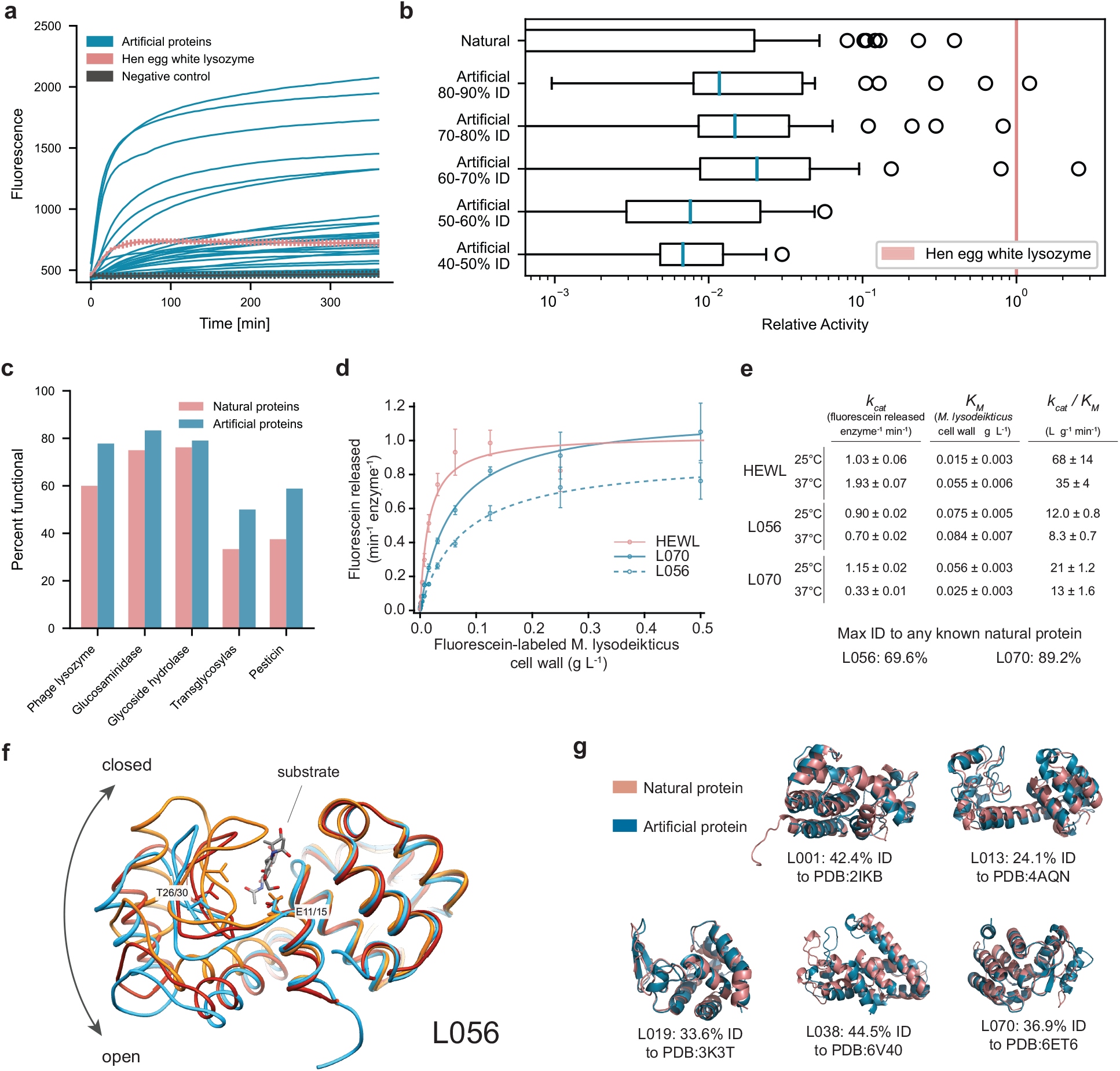
Artificial protein sequences are functional while reaching as low as 44% identity to any known protein, exhibit comparable catalytic efficiencies to a highly-evolved natural protein, and demonstrate similar structures to known natural folds. (a) Artificial proteins bind well to substrates and exhibit high fluorescence responses over time. Error bars (minimum and maximum) are shown for hen egg white lysozyme, HEWL, and negative (ubiquitin) controls. (b) Artificial proteins remain active even while being dissimilar (40-50% max ID i.e., top hit-identity) from known natural proteins. Outliers indicate high activity samples where relative activity is computed with respect to HEWL. (c) Artificial proteins are functional across protein families. Functional is defined as a fluorescence one standard deviation above the maximum value of all negative controls. (d) Michaelis-Menten kinetics of HEWL natural lysozyme (red) and two generated lysozymes (blue; L056 and L070) against cell-wall substrate show comparable performance (*n* = 3 technical replicates). (e) Michaelis-Menten constants of HEWL, L056, and L070 at different temperatures show that artificial proteins show comparable temperature stability. (f) We determined a 2.5 Å resolution crystal of L056 artificial lysozyme. A global overlay of L056 crystal structure with two representative T4 lysozyme conformations is shown with L056 presented in sky blue, ‘open’ conformation of M6I T4 lysozyme (PDB:150L) in dark red, ‘closed’ conformation of wild-type T4 lysozyme (PDB:3FA0) in orange, and substrate (PDB:148L) colored by element. Catalytic threonine (T30 in L056 and T26 in T4 lysozyme) and first catalytic glutamate (E15 in L056 and E11 in T4 lysozyme) are represented as sticks. (g) Although generated functional proteins are dissimilar from nature in sequence space, the predicted structures of generated proteins are similar to natural structures. To note, structural information was not provided in model training. Shown are predicted natural and generated structures with trRosetta.

In addition to a binary value for functionality, we calculated a relative activity score with respect to HEWL for the *in vitro* assay (see Methods). Our artificial proteins match activity levels of natural proteins even at lower levels of sequence identity to any known natural protein, (Fig 3b, Fig. S5). Notably a small number of proteins, both within the natural and artificial proteins, were within an order of magnitude of HEWL, which was substantially more active than all negative controls. These highly active outliers demonstrate the potential for our model to generate sequences that may rival natural proteins that have been highly optimized through evolutionary pressures.

From the 100 artificial proteins, we selected five proteins that spanned a wide range of max IDs (48-89%) to recombinantly express in *E. coli* (see Methods). Of these, only one, L008, generated no detectable expression (Fig. S6). Two (L013 and L038) expressed robustly to inclusion bodies and were not pursued further. Two proteins, L056 (max ID 69.6%) and L070 (max ID 89.2%) expressed well and incurred bactericidal activities towards the E. coli BL21(DE3) strain used during overnight induction at 16C. Spent media harbored enzymatic activity (see Methods), therefore, enzymes were purified from this material.

While both enzymes purified as monomers at the expected size by size exclusion chromatography, we also observed a defined later eluting (apparent lower molecular weight) species for each enzyme that corresponded to full-length enzyme by SDS-PAGE (Fig. S6). The K_M_ values of both monomers were too weak to be measured using a heterogeneous, fluorescein-labeled Micrococcus lysodeikticus cell wall substrate (Molecular Probes EnzChek Lysozyme kit); however, both monomers were active using a pseudo-first order kinetic assay (Fig. S7). In contrast, we could readily measure the K_M_ values for the purified apparent lower molecular weight species, where both L056 and L070 were highly active and had comparable Michaelis-Menten parameters to HEWL (Fig. 3e). Taken together, L056 and L070 harbor potent catalytic activity and bactericidal capabilities that are comparable to HEWL, while diverging from their nearest known natural sequence by 14 and 41 amino acids respectively, demonstrating that ProGen can generate artificial proteins with near native activity.

Next, we examined the structural divergence of the artificial proteins. We determined a 2.5 Å resolution crystal of L056 (see Methods, Figure 3f). The global fold was similar to predictions, with a Cα RMSD of 2.9 Å from the backbone structure predicted by trRosetta and 2.3 Å RMSD from a WT T4 Lysozyme structure^46,47^. The largest structural divergence occurs in the beta hairpin comprising residues 18-31. This region forms the bottom of the substrate binding cleft^48^ and is part of a hinge binding motion that is important for substrate binding^49^. The structure of the M6I mutant of T4 lysozyme (PDB 150L) is used as a model of the “open” state of this hinge and more closely resembles the structure of L056 (1.0Å Cα RMSD). Alignment with a structure featuring a covalently trapped substrate (PDB 148L) reveals that the active site cleft is well formed with the key catalytic residue Glu15 (Glu 11 in T4L) and key substrate binding residue Thr30 (Thr26 in T4L) correctly positioned. In addition, the hydrophobic core of L056 is well packed, with only two small packing voids of less than 5Å^3^ in volume, which is typical for structures of this size^50^. To additionally compare across structural representations, we utilized trRosetta^51^ to predict the structure of other functional artificial sequences. As in the crystal structure of L056, the predicted artificial structures roughly match known structures found in nature, Fig 3g. Collectively, the experimental crystal structure and the trRosetta calculations demonstrate that even though ProGen is not trained using any structural information or biophysical priors, it has likely learned structural invariances, presented as latent coevolutionary signatures, within large-scale evolutionary sequence databases. Furthermore, ProGen seems to capture these structural invariances in a generalizable fashion to design artificial proteins that reside within known natural protein folds while having high sequence dissimilarity, Fig 3g.

Trained on a universal protein sequence dataset spanning many families, ProGen designs proteins from any family when provided with the corresponding control tag as input. To explore this capability beyond the lysozyme clan, we evaluated ProGen’s performance in generating and predicting functional full-length sequences from families where other methods have previously been applied: chorismate mutase (CM)^6^ and malate dehydrogenase (MDH). We fine-tuned ProGen to unaligned protein sequences from these families and generated 64,000 unique sequences for each family (see Methods). Generated proteins exhibit similar conservation patterns to natural sequence libraries (Fig. 4ad). After aligning the generations to a sequence with known structure in Fig 4be, we observed that the conserved positions in generated sequences correlate with ligand-binding and buried residues. Using previously published sequences and their experimentally-measured assay data for CM^6^ and MDH^52^ proteins, we also evaluated the concordance of ProGen’s model likelihood for these sequences to their relative activity and compared it with the generative methods utilized in the original studies—bmDCA^6^ and proteinGAN^52^. Specifically, we measured per-token log-likelihoods for artificial sequences using ProGen (see Methods) and used them to predict if artificial sequences should function, which showed an area under the curve (AUC) of 0.85, significantly better (p<0.0001, two-tailed test, n = 1617) than bmDCA, which had an AUC of 0.78 (Fig. 4c). On MDH function data, ProGen log-likelihoods had an AUC of 0.94 (Fig. 4f), which was better than ProteinGAN discriminator scores, with an AUC of 0.87 (p<0.1, two-tailed test, n = 56). In sum, ProGen’s model likelihoods are better aligned with experimentally-measured assay data on two diverse protein datasets—chorismate mutase and malate dehydrogenase—than the sequence generation methods from original studies specifically tailored for these families.

**Figure 4:**
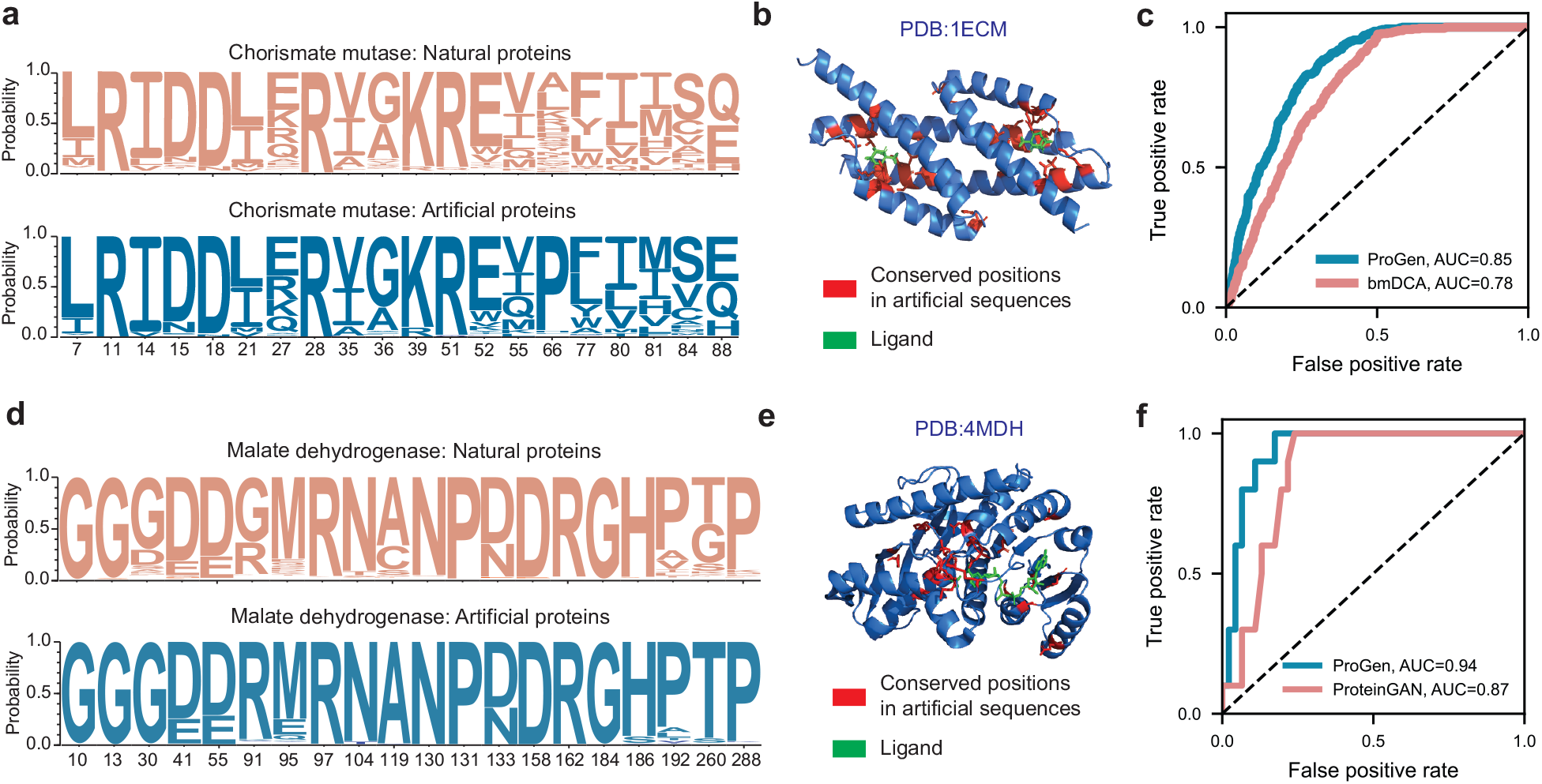
Applicability of conditional language modeling to other protein systems. Using the appropriate control tag, our language model, ProGen, can generate sequences for distinct protein families. Here we show that ProGen can generate chorismate mutase (CM) enzymes that exhibit a similar residue distribution to nature (a) and the conserved residues among generated sequences correlate to ligand-binding sites (b). ProGen’s model likelihoods can also accurately predict the functionality of CM variants from published data, slightly better than the coevolutionary bmDCA^6^ algorithm from the original study (c). ProGen can also generate malate dehydrogenase (MDH) proteins that exhibit a similar residue distribution to nature (d). The conserved residues among generated sequences correlate to buried residues (e). ProGen’s model likelihoods are also accurate in predicting functionality of published variants of MDH, similar to the generative proteinGAN^52^ model used in the original study (f).

To understand the relative impact of the universal sequence dataset and of sequences from the protein family of interest on ProGen’s generation ability, we perform two ablation studies using the CM and MDH experimentally-measured assay data. First, we evaluated the performance of ProGen trained only with the universal sequence dataset. We measured per-token log-likelihoods for artificial sequences using this version of ProGen using control tags for CM and MDH (see Methods). These likelihoods showed in a significant drop in AUC of 0.18 for CM (p<0.0001, two-tailed test, n = 1617) and 0.08 for MDH (p<0.1, two-tailed test, n = 56), as compared to fine-tuned ProGen when predicting if an artificial sequence should function. Conversely, the ProGen architecture trained on CM and MDH protein sequences alone without the benefit of initial training on the universal sequence dataset also showed a significant drop in performance as compared to fine-tuned ProGen—the AUC reduced by 0.11 (p<0.0001, two-tailed test, n = 1617) and 0.04 (p<0.05, two-tailed test, n = 56) on the CM and MDH data respectively. These results indicate that both components of our training strategy—initial training on the universal sequence dataset and fine-tuning on the protein family of interest—contribute significantly to final model performance. Training with the universal sequence dataset containing many protein families enables ProGen to learn a generic and transferable sequence representation that encodes intrinsic biological properties. Fine-tuning on the protein family of interest steers this representation to improve generation quality in the local sequence neighborhood. Similar to the adaptability shown by neural networks trained on large datasets using transfer learning and fine-tuning in natural language processing^20,53,54^ and computer vision^37,55^, protein language models have the potential to be a versatile tool for generating tailored proteins with desired properties.

In conclusion, our study shows that a state-of-the-art transformer-based conditional language model trained only with evolutionary sequence data generates functional artificial proteins across protein families. Additional analyses suggest that our model has learned a flexible protein sequence representation which can be applied to diverse families such as lysozymes, chorismate mutase, and malate dehydrogenase. Applications of our model could include generating synthetic libraries of highly-likely functional proteins for novel discovery or iterative optimization. In combination with ever-increasing sources of sequence data and more expressive control tags, our work points to an exciting potential for the use of deep learning-based language models for precise design of novel proteins to solve important problems in biology, medicine, and the environment.

## Methods

### Training Data Curation

The initial training data consisted of over 281 million non-redundant, annotated protein sequences with associated metadata, formulated as control tags. The amino acid vocabulary consisted of the standard 25 amino acids designations in IUPAC^56^. The control tags were divided into two categories: (1) keyword tags and (2) taxonomic tags. Following the definitions laid out in the UniprotKB controlled, hierarchical vocabulary of keywords (many of which are derived from Gene Ontology (GO) terms^57^), the control keyword tags included 1100 terms ranging from cellular component, biological process, and molecular function terms. The taxonomic tags include 100k terms from the NCBI taxonomy across the eight standard taxonomic ranks. The aggregated dataset was split into a training set of size 280 million and two test sets, an out-of-distribution test set (OOD-test) of size 100k from 20 protein families and a randomly sampled in-domain test set (ID-test) of size 1 million, that were held-out for training and used for evaluation. After model training on the training database, the model was further trained, i.e. fine-tuned, to the following datasets for generation and classification tasks.

For fine-tuning on lysozyme proteins, five protein families from the Pfam database were selected, Phage lysozyme (PF00959), Pesticin (PF16754), Glucosaminidase (PF01832), Glycoside hydrolase family 108 (PF05838), and Transglycosylase (PF06737), yielding a total of 55,948 sequences. Proteins were provided to the model during fine-tuning as unaligned protein sequences with one control tag prepended for the protein family designation. For fine-tuning on chorismate mutase proteins, a search with HHBlits and blastp was performed with residues 1-95 of EcCM (the CM domain of the E. coli CM-prephenate dehydratase, the P-protein) yielding 20214 sequences. For fine-tuning on malate dehydrogenase proteins, the L-lactate/malate dehydrogenase protein family from Interpro IPR001557 was selected with 17094 sequences.

### Conditional Language Modeling

Our conditional language model, named ProGen, is a decoder style Transformer^19^-variant with 36 layers, 8 attention heads per layer, and dimensions *d* = 1028 and *f* = 512. To learn the conditional distributions over amino acids and control tags, a sequence containing n tokens is embedded as a sequence of *n* corresponding vectors in ℝ^*d*^. This sequence of vectors is stacked into a matrix *X*_0_ ∈ ℝ ^*n×d*^, and is processed by a series of *l* pre-activation layer-normalized Transformer self-attention layers^23^. Logits for each token in the vocabulary are computed via a learned linear transformation that is applied to each output of the last layer. During training, these logits input to a cross-entropy loss. During generation, the logits for the final token in the sequence are normalized with a softmax to get a distribution for sampling a new token.

### ProGen Training

For training, we included each sequence and its reverse. We prepended each sequence with a corresponding subset of control tags. For a given sequence, there can be multiple versions across databases, each with their own associated control tags. We randomly sampled which set of control tags to utilize, but biased sampling toward SwissProt tags as they are manually verified. Additionally, we always included a sample with the sequence alone without control tags so that ProGen could be used to complete proteins using sequence data alone. We truncated all sequences to a maximum length of 512. Sequences of length less than 512 were padded, and padded tokens were excluded from the cost function used for training. Our model was implemented in TensorFlow and trained with a global batch size of 2048 distributed across 256 cores of a Cloud TPU v3 Pod for 1 million iterations. Training took approximately two weeks using Adagrad with linear warmup from 0 to 1e^-2^ over the initial 40k steps with a linear decay for the remainder of training. The model was initialized with pretrained weights of CTRL^23^, which was trained on an English language corpus.

### Lysozyme Generation

We fine-tuned ProGen to the 55,948-sequence fine-tuning dataset using the conditional language modeling loss function introduced in Eq. 1, using a separate control tag for each of the five lysozyme families. The model was fit for 4 epochs using the Adam optimizer^58^ with a learning rate of 0.0001, batch size of 2, gradient norm clipping^59^ threshold of 0.25, and a dropout^60^ rate of 0.1. We then applied sampling using the final checkpoint of the fine-tuned model. We generated one million de novo sequences from the learned conditional probability distribution *p*_*θ*_ (*a*_*i*_|*a*_<*i*_, *c*) using each of the 5 lysozyme families as a control tag *c*, and applying top-p sampling^61^, which zeros out the probability of the tail of the distribution during sampling, and uses a hyperparameter *p* to determine what fraction of the original distribution to keep. Lower *p* values result in sequences with a higher likelihood under the model, but lower diversity. We generated a batch of one million synthetic sequences (Fig. S3) using *p* values that varied in [0.25,0.50,0.75], and applied the sequence selection criteria in the next section to determine which sequences to synthesize.

### Lysozyme Sequence Selection

We selected sequences for synthesis by ranking them using the combination of an adversarial discriminator^22,43^ and generative model log-likelihood scoring^44^. First, we trained an adversarial discriminator to distinguish between natural lysozymes and ProGen-generated lysozymes. A higher discriminator score indicates a protein sequence that is “semantically” and “grammatically” closer to natural sequences, but not necessarily one of high sequence identity to natural proteins. To train the discriminator, we generated a batch of samples from fine-tuned ProGen (with nucleus sampling turned off, or *p* = 1) that was the same size and distribution of families as our dataset of natural lysozymes. The discriminator architecture was a fine-tuned TAPE-BERT^34^. For robustness, we trained three discriminators using different random seeds. We assigned each sequence a discriminator score as the geometric mean of the probability of the sample being a natural sequence as predicted by the three discriminators. We also assigned each sequence a log-likelihood score as the average per-token log-likelihood for each sample computed using the fine-tuned ProGen model and conditioned on the control tag used to generate the sequence, given by

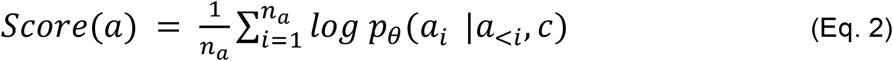

A higher log-likelihood score indicates a sequence close to the probability distribution of sequences seen in training. We selected proteins for de novo sequences using separate rankings based on the discriminator and log-likelihood scores. We separately ranked candidate sequences in maximum sequence identity ranges of 40-50%, 50-60%, 60-70%, 70-80%, and 80-90%. For each range, we added the top discriminator-ranked sequences, skipping any sequences that were >80% identical to any previously selected sequence, for a total of 90 sequences. Ten more sequences were added based on ranking by generative model log-likelihood scores in each range, again skipping any sequences with >80% identity to any previously selected sequence.

### Evaluating ProGen on Other Protein Systems

We also evaluated ProGen on generation of Chorismate Mutase (CM) and Malate Dehydrogenase (MDH) proteins. We separately fine-tuned ProGen on datasets of CM and MDH proteins using the Adam optimizer, a learning rate of 1e-4, a gradient norm clipping threshold of 0.25, and a dropout rate of 0.1. We also preappended the CM and MDH data with control tags that corresponded to CM and MDH families in ProGen’s original training. After fine-tuning, we generated a set of 64k sequences using top-p sampling (*p* = 0.75) from the CM and MDH fine-tuned models respectively. We measured concordance of our model’s log-likelihoods with protein function data on CM and MDH sequences, and compared with bmDCA^6^ and ProteinGAN^52^ baselines respectively. We computed the area under the curve (AUC) in receiver operating characteristic (ROC) curves for predicting binary function labels from model scores. We computed model scores for each sequence in both CM and MDH by using the per token model log-likelihood in Eq. 2. We used model scores for bmDCA given by negative energy of each CM sequence provided by the study’s authors^6^. We also applied thresholding at 0.42 norm relative enrichment (r.e.) to obtain binary labels for CM function, which roughly corresponds to the cut-off point between two modes that exist in CM function data, to be used for ROC curves, following the original study^6^.

Since model likelihoods for GANs are intractable, we used discriminator scores corresponding to the probability at which the ProteinGAN discriminator predicted each sample was real as a ProteinGAN model score for each MDH sequence. The MDH functional labels are binary, so no thresholding was needed to compute AUCs. For an ablation study on ProGen, we also evaluate 1. a randomly initialized LM that has the same architecture as ProGen and is finetuned to the same task-specific data as ProGen (CM or MDH), but is not pretrained on a larger dataset 2. ProGen without task specific fine-tuning, conditioning on control tags for CM or MDH from ProGen’s original pretraining data. After measuring the AUC of each model for each dataset, we used bootstrapping to compute the statistical significance of the difference in AUC of fine-tuned ProGen vs. the reference method (bmDCA and ProGen ablations for CM, ProteinGAN and ProGen ablations for MDH). At each bootstrapping iteration, we resampled a new dataset of fitness and model score pairs the same size as the original dataset by randomly selecting data points from the original dataset with replacement. For each sample dataset, we compute the difference in AUC score between fine-tuned ProGen and the reference method. We drew a total of 10,000 bootstrapping samples, and the p-value is given by the percentage of the samples where the baseline achieves an AUC greater than or equal to fine-tuned ProGen, multiplied by two to give two-tailed.

### Materials

All reagents were purchased from Thermo Fisher Scientific (Waltham, MA, USA) unless otherwise noted. DNAs utilized for in vitro translation were purchased from Twist Bioscience (San Francisco, CA, USA) and DNAs utilized for *E. coli* expression and purification were purchased from VectorBuilder (Chicago, IL, USA).

### High-throughput cell-free expression of lysozymes

Lysozymes were expressed using the Tierra Bioscience (San Leandro, CA) cell-free expression platform. Cell-free extracts for protein expression were prepared according to the methods of Sun *et al*. (2013)^62^ with the following modifications: Terrific Broth was used in lieu of 2xYT, cells were lysed in a single pass by French® press at 10,000 PSI, dithiothreitol was omitted from wash buffers, and run-off and dialysis steps were removed to streamline extract processing. Expression reactions were composed of cell-free extract, an energy buffer, and a linear DNA template containing a promoter sequence, the protein sequence of interest, the sequence of a strep purification tag and a terminator sequence; reactions were carried out at 29 °C for 6 hours. Expression reactions for screening optimal affinity purification tag terminus were performed in 10µL volumes; selected reactions with good expression were then scaled to 200µL. Lysozymes were purified from expression reactions by affinity chromatography with elution by enzymatic cleavage with 3C protease leaving a small sequence scar.

### High-throughput screening of lysozyme activity

Purified cell-free synthesized lysozymes were assayed with the EnzChek® Lysozyme Assay Kit (Thermo Fisher Scientific, Waltham, MA). The assay was performed according to protocol with minimal modifications. Hen egg white (HEW) lysozyme standards and purified proteins in buffer (100mM Tris pH 7.4, 150mM NaCl, 2mM TCEP, 20% glycerol) were brought to 50µL with reaction buffer (100mM sodium phosphate pH 7.5, 100mM NaCl, 2mM NaN_3_) in a 96 well plate. 50µL of DQ lysozyme substrate, fluorescein conjugate (1 mg/mL) was added to each well and fluorescence (ex 485/20; em 528/20) was collected every 5 minutes with a Synergy 2 multi-mode microplate reader (BioTek, Winooski, VT) for 6 hours at 37 °C.

For each 96-well plate, three random wells were dedicated for egg white lysozyme controls and three wells were dedicated for a negative control of ubiquitin expressed and purified on the Tierra Biosciences cell-free expression platform. A purified protein was considered functional if it exhibited a higher fluorescence than one standard deviation above the maximum fluorescence value of all negative controls. The relative activity for each protein was calculated by the following equation:

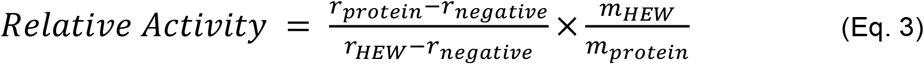

Where r is the linear rate of fluorescence increase in the initial twenty minutes of the fluorogenic assay and m is the mass of protein as determined by Bradford assay concentration and measured volumes.

### *E. coli* Expression of Lysozyme Variants

We chose five generated lysozyme variants (L008, L013, L038, L056, L070) for expression in *E. coli* based on strength of signal in the in vitro assay, expression level in the cell-free system, and max ID to natural proteins. Generated lysozyme variants, were codon optimized for *E. coli* (Integrated DNA Technologies) with an HRV3C protease site N-terminal of the open reading frame. DNA was synthesized and cloned in-frame with a 5’ His_6_-tag in a pET vector and transformed into BL21(DE3) (Vectorbuilder). One liter of Terrific Broth (Fisher) was pre-warmed to 37°C before being inoculated with 10mL of an overnight starter culture. Cultures were grown to 0.6 < OD_600_ < 1.0 before temperature was dropped to 16°C for expression. Cultures were induced with 0.5 mM IPTG (source) and protein expression allowed to continue overnight. For induced cultures of L056 and L070, significant turbidity was observed in the spent media after cells were pelleted at 3,500 r.c.f. for 30 mins at 4°C. Spent media also harbored lysozyme activity as ascertained through fluorescence increase over time of the Fluorescein-labeled *M. Lysodeikticus* cell wall substrate (EnzChek kit; ThermoFisher). Spent media was saved for protein purification (outlined below) and cell pellet frozen and stored at - 20°C. Variant L008 did not express under multiple different conditions. L013 and L038 expressed highly to inclusion bodies.

### Purification of L056 and L070 from spent media

Media was split into two 0.5 L pools each. The first pools were loaded onto a 5 mL HisTrap FF NiNTA column (GE) using a peristaltic pump at room temperature. Columns were washed with 200 mL of 30 mM HEPES pH 7.6, 150mM NaCl, 25 mM imidazole, 0.5 mM TCEP. Columns were eluted with 25 mL of 30 mM HEPES pH 7.6, 150mM NaCl, 250 mM imidazole, 0.5 mM TCEP. Eluates were concentrated to 8-10mL and dialyzed against 30 mM HEPES pH 7.6, 150mM NaCl, 0.5 mM TCEP with HRV3C protease added overnight at 4°C. Dialyzed protein was put through an ortho 5 mL HisTrap FF NiNTA column (GE) to remove HRV3C protease and uncleaved lysozyme. Though highly pure by SDS-PAGE analysis, protein was further purified by size-exclusion chromatography and loaded on an S75 10/300 gl column pre-equilibrated with 30 mM HEPES pH 7.6, 150mM NaCl, 0.5 mM TCEP. Two peaks were resolved for each variant and that harbored lysozyme activity against the fluorescein-labeled *M. Lysodeikticus* cell wall substrate (EnzChek kit; ThermoFisher). Individual peaks were pooled and protein concentration determined either by Bradford assay (Biorad) or by SDS-PAGE using colloidal coomassie (ThermoFisher) and HEWL in-gel standards.

The second spent media pools were batch-bound to 5 mL of HisPur NiNTA resin (ThermoFisher) at 4°C for 1 hour before protein-bound resin was pelleted through centrifugation at 3000 r.c.f. for 5 mins at 4°C. Protein bound resin was resuspended with 25 mL 30 mM HEPES pH 7.6, 150mM NaCl, 25 mM imidazole, 0.5 mM TCEP and applied to a gravity flow column (BioRad) at room temperature. Columns were washed with 200 mL of 30 mM HEPES pH 7.6, 150mM NaCl, 25 mM imidazole, 0.5 mM TCEP. Columns were eluted with 25 mL of 30 mM HEPES pH 7.6, 150mM NaCl, 250 mM imidazole, 0.5 mM TCEP. Eluates were concentrated to 8-10mL and dialyzed against 30 mM HEPES pH 7.6, 150mM NaCl, 0.5 mM TCEP with HRV3C protease added overnight at 4°C. Lysozyme was separated from HRV3C protease by size-exclusion chromatography on an S75 10/300 gl column pre-equilibrated with 30 mM HEPES pH 7.6, 150mM NaCl, 0.5 mM TCEP. Two peaks were resolved for each variant and that harbored lysozyme activity against the fluorescein-labeled *M. Lysodeikticus* cell wall substrate (EnzChek kit; ThermoFisher) that corresponded to peaks observed in the first pool purification. Individual peaks were pooled and protein concentration determined either by Bradford assay (Biorad) or by SDS-PAGE using colloidal coomassie (ThermoFisher) and HEWL in-gel standards.

### Michaelis-Menten kinetics of lysozyme variants using fluorescein-labeled *M. lysodeikticus* cell wall

Fluorescein-labeled *M. Lysodeikticus* cell wall substrate (EnzChek kit; ThermoFisher) was reconstituted in 30mM HEPES pH 7.6, 150mM NaCl to 1 mg/mL, aliquoted and stored at -20°C until use. A serial two-fold dilution series of substrate was prepared in 30mM HEPES pH 7.6, 150mM NaCl and treated as a 2X solution for enzymatic assays. Enzyme concentration was calculated either through Bradford assay (Bio-Rad) or by SDS-PAGE, in-gel using Novex Colloidal Coomassie stain against a hen egg white lysozyme (HEWL; Alfa Aesar) standard. Enzymes were diluted to between 10 and 100nM in 30mM HEPES pH 7.6, 150mM NaCl (HEWL) or 30mM HEPES pH 7.6, 150mM NaCl, 0.5mM TCEP (L056 and L070) and these stocks treated as a 2X solution for enzymatic assays. Kinetic assays were performed in a Tecan Spark 10M plate reader using monochrometers with a fixed 20 nm bandpass filter in a 384-well black-bottom plate (Corning) at 10 µL final volume. Reactions were initiated by pipetting 5 µL of substrate into appropriate wells followed immediately by 5 µL of enzyme, mixed by pipetting before starting data acquisition. The dead-time from reaction initiation to acquisition of first read was measured to be 24 s. For reactions carried out above ambient temperature (25°C), the plate was preincubated at temperature for at least 5 mins prior to reaction initiation. Initial velocities were calculated through linearly fitting fluorescence intensity (au) versus time for the first 2 minutes of each reaction. Finally, velocities were converted from au to fluorescein liberated through application of a fluorescein (Sigma) standard curve (Figure S6) and normalized to enzyme concentration. Averaged data (*n* = 3 technical replicates) were nonlinearly fit to the Michaelis Menten model (Eq. 4) in IgorPro 7 to report *k*_*cat*_ in units of fluorescein liberated enzyme^-1^ min^-1^ and *K*_*M*_ in units of g L^-1^ (the average molecular weight of the fluorescein-labeled *M. Lysodeikticus* cell wall substrate was unknown and likely heterogeneous).

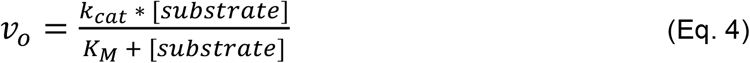

### Lysozyme *k*_*cat/*_*K*_*M*_ extrapolation from pseudo first-order kinetic data

For the higher molecular weight L056 and L070 species whose *K*_*M*_’s were beyond the concentration regime of the fluorescein-labeled *M. Lysodeikticus* cell wall substrate (EnzChek kit; ThermoFisher), the ration kcat/KM was measured through pseudo first-order kinetics where when [Enyzme] >> Substrate the Michaelis-Menten model simplifies to Eq. 5. Fluorescein-labeled *M. Lysodeikticus* cell wall substrate was diluted to 0.01 g L^-1^ and this stock was treated as 2X for kinetic assays. Kinetic assays were performed in a Tecan Spark 10M plate reader using monochrometers with a fixed 20 nm bandpass filter in a 384-well black-bottom plate (Corning) at 10 µL final volume. The dead-time from reaction initiation to acquisition of first read was measured at 24 s and the 0 s fluorescence intensity was measured through dilution of substrate with buffer. Reactions were initiated by pipetting 5 µL of 2X enzyme into 5 µL of 2X substrate in a prewarmed 384-well black assay plate (Corning). Five technical replicates were performed across four enzyme concentrations. The resultant data were not described by a single exponential model but were described by a double exponential model (Eq. 6), likely due to the heterogeneity of the substrate, and all data were fit in IgorPro7. The reciprocal of the weighted sum of each tau component was taken to estimate a single *k*_*obs*_ value for subsequent analysis (Eq. 7). To estimate *k*_*cat*_*/K*_*M*_, *k*_*obs*_ values were plotted against enzyme concentration where the slope of a linear fitting is equal to *k*_*cat*_*/K*_*M*_.

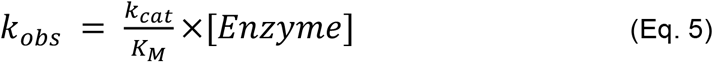

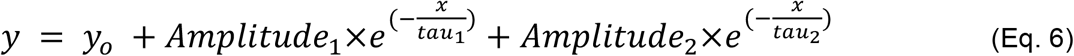

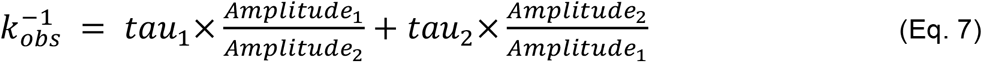

### Crystallization and structure determination of L056

Purified L056 was concentrated to 18.6 mg/mL in 30 mM HEPES pH 7.6, 150 mM NaCl, 0.5 mM TCEP. Crystals were identified from sitting drop vapor diffusion experiments set at 20°C with a 1:1 ratio of 200nL protein and 200nL well solution (0.1 M CHES 9.5 pH, 30 %w/v PEG 3K). Diffraction data were collected from a single crystal at Beamline 8.3.1. at the Advanced Light Source. Data were processed using XDS^63^ and a molecular replacement solution was identified using phaser^64^ with a trRosetta model of L056 as a search model. Significant translational non-crystallographic symmetry and differences with the search model resulted in maps that were initially hard to interpret. The initial model was improved using Refmac jelly body refinement^65^ prior to rebuilding using phenix.autobuild^66^ and the CCP4 buccaneer_pipeline^67^. The model was finalized and iteratively improved with multiple rounds of manual modification in Coot^68^ and refinement using phenix.refine^69^. The model is deposited as PDB 7RGR.

## Supporting information

Structure Validation Report

## Author Contributions

A.M. conceived and designed the study in collaboration with S.S. A.M. and B.K. designed and performed machine learning modeling, generation, and scoring. B.P.M. performed the cell-free expression and activity assay and was supervised by Z.Z.S. E.G. performed the cell-based expression and kinetics assay and was supervised by J.S.F. J.H., J.L.O., J.S.F. performed the structure determination. A.M., S.S., B.K., and N.N. performed computational analysis, and were advised by C.X. R.S. provided advice on machine learning and computational methods. A.M., J.S.F., and N.N. wrote the manuscript with feedback and contributions from all authors, in particular from E.G. and B.K. N.N. supervised and managed the project.

## Acknowledgements

We thank Bryan McCann, Christine Gee, Eric Procko, Kelly Trego, Lav Varshney, Nitish Shirish Keskar, and Silvio Savarese for their feedback at various stages of this project. We thank Audrey Cook, Denis Lo, and Vanita Nemali for operational support. Thanks also to the Salesforce Research Computing Infrastructure team and the Google Cloud TPU team for their help with computing resources, in addition to Twist Bioscience for DNA synthesis support. Beamline 8.3.1 at the Advanced Light Source is operated by the University of California Office of the President, Multicampus Research Programs and Initiatives grant MR-15-328599, the National Institutes of Health (R01 GM124149 and P30 GM124169), Plexxikon Inc., and the Integrated Diffraction Analysis Technologies program of the US Department of Energy Office of Biological and Environmental Research. The Advanced Light Source (Berkeley, CA) is a national user facility operated by Lawrence Berkeley National Laboratory on behalf of the US Department of Energy under contract number DE-AC02-05CH11231, Office of Basic Energy Sciences. JSF was supported by NIH GM123159 and a Sanghvi-Agarwal Innovation Award.

## Supplementary Materials

**Tables S1-S5**

**Figures S1-S7**

