## Supplemental Information for "Deep neural language modeling enables functional protein generation across families"

### Supplementary Information for:
### Deep neural language modeling enables functional protein generation across families

Ali Madani[1*], Ben Krause[1], Eric R. Greene[2], Subu Subramanian[3,4], Benjamin P. Mohr[5], James M. Holton[6,7,8], Jose Luis Olmos, Jr.[2], Caiming Xiong[1], Zachary Z. Sun[5], Richard Socher[1], James S. Fraser[2], Nikhil Naik[1*]

1. Salesforce Research, Palo Alto CA 94301, USA.
2. Department of Bioengineering and Therapeutic Sciences, University of California, San Francisco, San Francisco, CA 94158, USA.
3. Department of Molecular and Cell Biology, University of California, Berkeley, Berkeley, CA 94720, USA.
4. Howard Hughes Medical Institute, University of California, Berkeley, Berkeley, CA 94720, USA.
5. Tierra Biosciences, San Leandro, CA 94577, USA.
6. Molecular Biophysics and Integrated Bioimaging Division, Lawrence Berkeley National Laboratory, Berkeley, CA 94720, USA.
7. Stanford Synchrotron Radiation Lightsource, SLAC National Accelerator Laboratory, Menlo Park, CA 94025, USA.
8. Department of Biochemistry and Biophysics, University of California, San Francisco, San Francisco, CA 94158, USA.

**Contents of this file include:**

Tables S1-S5
Figures S1-S7
Supplementary References

| | Dataset name | Dataset size used | Purpose for model training |
|---|---|---|---|
| (a) | Swiss-prot | 400k protein sequences, 1000 unique UniprotKB keywords | High-fidelity sequences + metadata for training |
| (b) | TrEMBL | 180M protein sequences, 20 unique UniprotKB keywords | Higher-quantity, lower-fidelity proteins for training |
| (c) | UniParc | 280M protein sequences | Reference database used for full-range of proteins exposed to model |
| (d) | NCBI Taxonomy | 100k unique taxonomy terms | Provide source organism information to the model |
| (e) | Pfam | 56k protein sequences | Curated protein families for lysozymes |
| (f) | Uniref30 | 7k protein sequences | Chorismate mutase sequences used from an HHBlits search |
| (g) | NCBI nr database | 13k protein sequences | Chorismate mutase sequences used from a blastp search |
| (h) | Interpro | 17k protein sequences | Malate dehydrogenase proteins under IPR001557 |

**Table S1:** A list of publicly available datasets used to train and fine-tune ProGen. The training datasets (a-c) contain a total of 280 million unique proteins which are associated with 101,100 tags based on keywords related to biological processes and molecular function and taxonomic information (d). The fine-tuning datasets (e-h) are used to further improve ProGen's ability to generate protein sequences in local sequence neighborhoods of lysozymes, chorismate mutase, and malate dehydrogenase proteins.

**Template: <$c_1$> <$c_2$> . . . <$c_N$> $a_1$ $a_2$ $a_3$ $a_4$ $a_5$ . . .**

**Training Sample**

```
<Metazoa><Chordata><Mammalia><Rodentia><Muridae><Rattus><Rattus><Norvegicus>
<NAD>
<Translocase>
<Iron><Iron-sulfur><2Fe-2S>
<Mitochondrion><Mitochondrion inner membrane>
<Transport><Electron transport><Respiratory chain>
MFSLALRARASGLTAQWGRHARNLHKTAVQNGAGGALFVHRDTPENNPDTPFDFTPENYERIEAIVRNYPEGHRAA
AVLPVLDLAQRQNGWLPISAMNKVAEVLQVPPMRVYEVATFYTMYNRKPVGKYHIQVCTTTPCMLRDSDSILETLQ
RKLGIKVGETTPDKLFTLIEVECLGACVNAPMVQINDDYYEDLTPKDIEEIIDELRAGKVPKPGPRSGRFCCEPAG
GLTSLTEPPKGPGFGVQAGL
```

**Fine-tuning Sample**

```
<Phage Lysozyme>
MNIFEMLRIDEGLRLKIYKDTEGYYTIGIGHLLTKSPSLNAAKSELDKAIGRNCNGVITKDEAEKLFNQDVDAAV
RGILRNAKLKPVYDSLDAVRRCALINMVFQMGETGVAGFTNSLRMLQQKRWDEAAVNLAKSRWYNQTPNRAKRVI
TTFRTGTWDAYKNL
```

**Figure S1:** Sample sequences with associated control tags provided for model training during training and fine-tuning. The amino acid sequence $(a_1, a_2, \ldots)$, prepended with desired control tags ($< c_1 >, < c_2 >, . . .$), are formulated as tokens for ProGen to autoregressively compute the probability of the next token. Control tags are constructed to provide information regarding molecular function, biological process, cellular component, or curated protein family.

| Pfam name (ID) | Description | Number of sequences in fine-tuning set | Average sequence length in fine-tuning set | Number of artificial sequences tested | Average sequence length of artificial sequences tested |
|---|---|---|---|---|---|
| Phage lysozyme (PF00959) | Glycoside hydrolase family 24 which includes lambda phage lysozyme and Escherichia coli endolysin | 16488 | 151 (±29) | 20 | 170 (±16) |
| Glyco_hydro_108 (PF05838) | Glycoside hydrolase family 108 | 5857 | 152 (±38) | 45 | 179 (±6) |
| Glucosaminidase (PF01832) | Glycoside hydrolase family 74 | 23238 | 138 (±14) | 8 | 139 (±4) |
| Transglycosylase (PF06737) | Resuscitation-promoting factor proteins | 9824 | 84 (±7) | 8 | 93 (±16) |
| Pesticin (PF16754) | Hydrolase enzyme secreted by Yersinia pestis and other Gammaproteobacteria to kill related bacteria occupying ecological niche. | 541 | 167 (±28) | 19 | 176 (±17) |

**Table S2:** The five lysozyme families utilized for *de novo* protein sequence generation, along with details of the fine-tuning dataset. We utilize a total of 55,948 sequences from these five families obtained from Pfam, ranging from 541 to 23238 proteins per family. The proteins within each family exhibit a range of average sequence lengths, from 84 to 167 residues (one standard deviation shown in table), and ProGen is able to generate similarly diverse sequence average length statistics, from 93 to 179 residues.

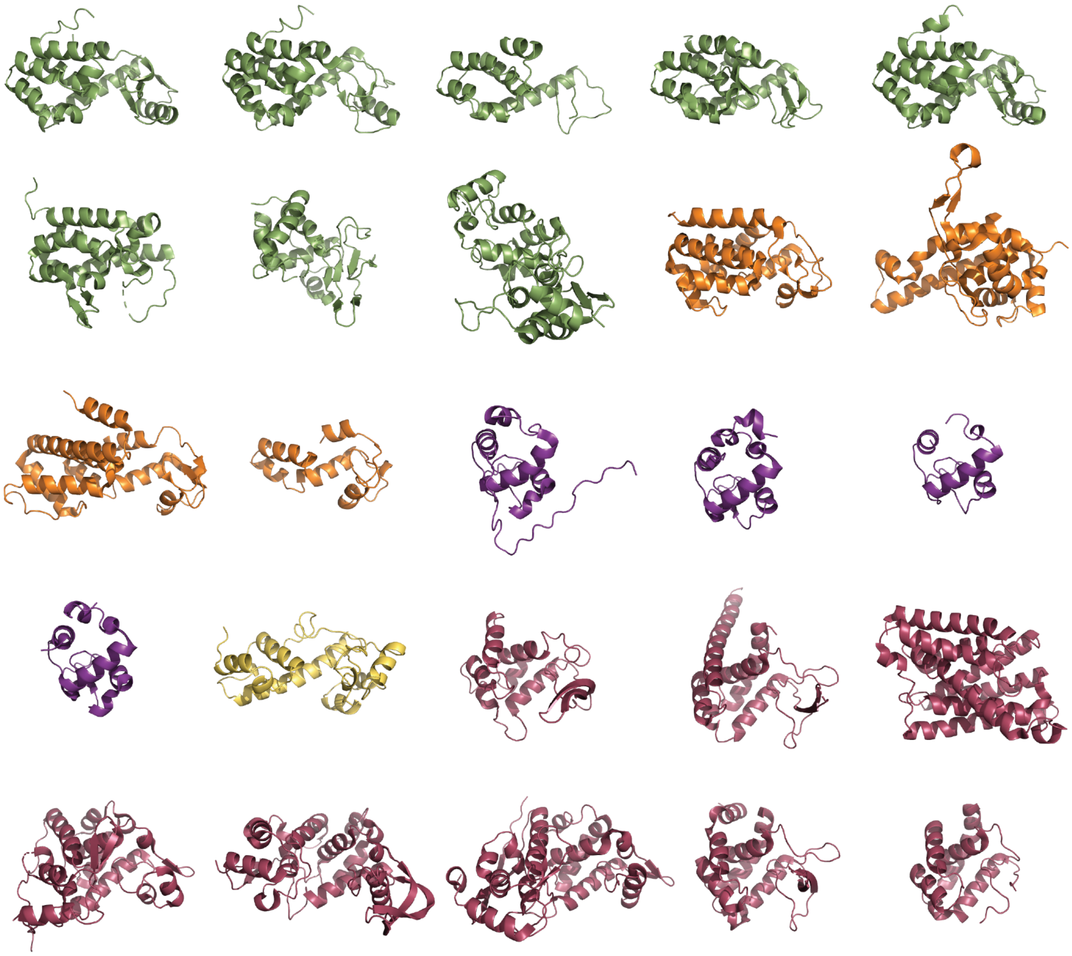

**Figure S2:** Crystal structures of known natural proteins within the five lysozyme families selected in this study, where Phage lysozyme, (PF00959), Glyco_hydro_108 (PF05838) Transglycosylas (PF06737), Pesticin (PF16754), and Glucosaminidase (PF01832) families (with Pfam IDs) are represented as green, orange, purple, yellow, red respectively. The five families exhibit considerable structural diversity and multiple structural folds, presenting a challenging design space for a sequence-only model.

| | Diversity-50% [number of clusters] | Diversity-80% [number of clusters] |
|---|---|---|
| **Natural lysozyme** | 1021 | 4115 |
| **Artificial lysozyme** | 1193 | 6206 |
| **Natural chorismate mutase** | 831 | 1324 |
| **Natural malate dehydrogenase** | 343 | 1231 |

**Table S3:** Sequence diversity of natural lysozymes, ProGen-artificial lysozymes, natural chorismate mutase, and natural malate dehydrogenase proteins. The sequence statistics indicate that lysozymes are more sequence-diverse than the other two protein systems. Also, ProGen is able to generate artificial sequences that exhibit higher diversity than its corresponding natural lysozyme library. Each sequence database is clustered by mmseqs2 with 50%/80% max identity with 80% coverage. The value of the diversity metric is the number of clusters with three or more members. A higher value represents a more diverse library.

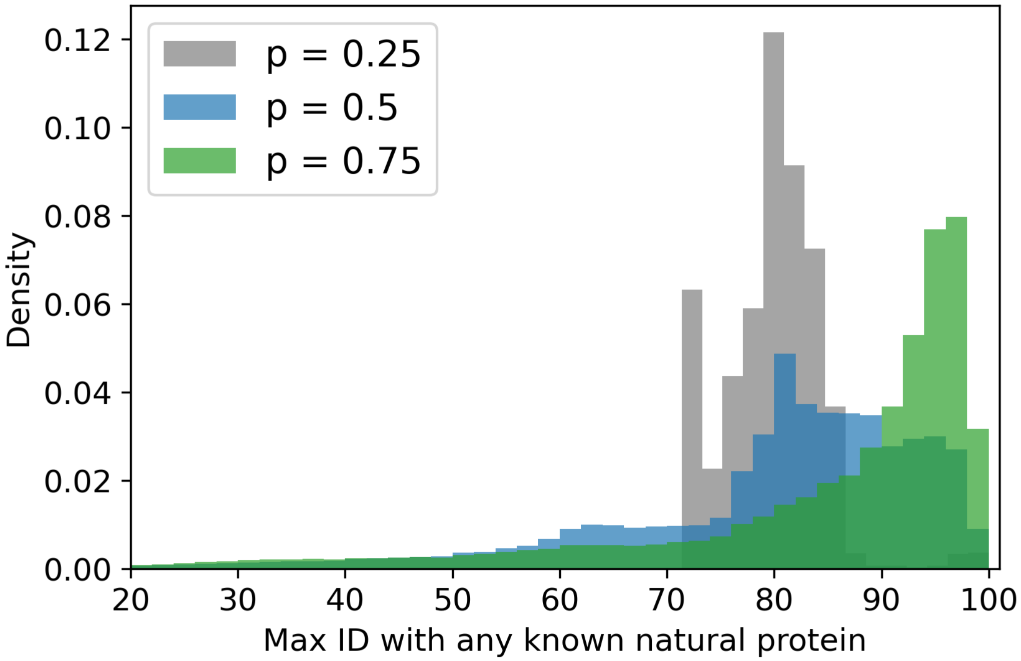

**Figure S3:** A density plot of max identity distribution of one million lysozyme sequences generated by ProGen at different top-p values, which is a hyperparameter used for controlling the diversity of generated sequences. A lower p setting allows the model to generate sequences that have a higher likelihood under the model but lower diversity, meaning that the mean max identity of the generated sequences may be closer to 100, but the spread is smaller. In contrast, a higher p setting results in a more diverse distribution, but at the cost of more potential mistakes. In our experiments, we generate a total of one million sequences at three different top-p settings and select a hundred from this pool for expression and characterization.

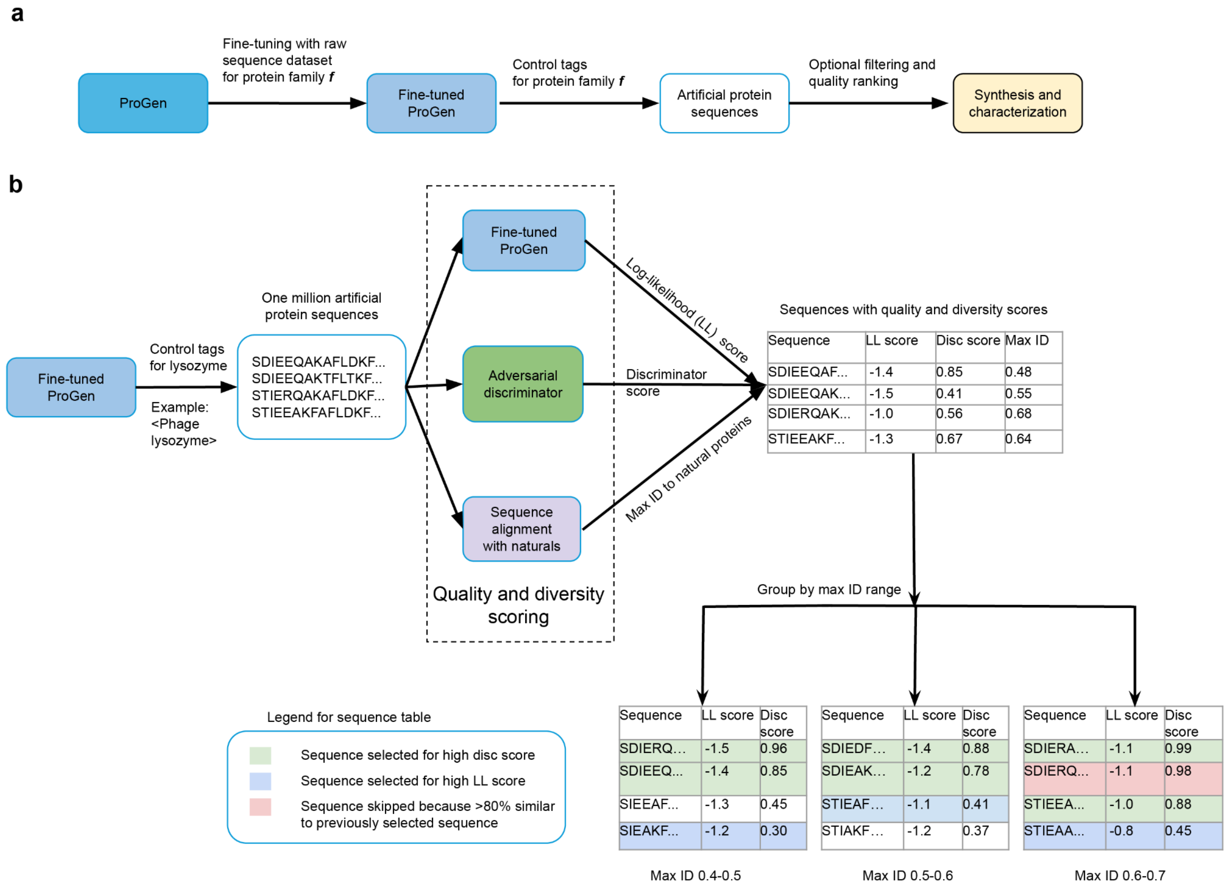

| Sequence | LL score | Disc score | Max ID |
|---|---|---|---|
| SDIEEQAF... | -1.4 | 0.85 | 0.48 |
| SDIEEQAK... | -1.5 | 0.41 | 0.55 |
| SDIERQAK... | -1.0 | 0.56 | 0.68 |
| STIEEAKF... | -1.3 | 0.67 | 0.64 |

| Sequence | LL score | Disc score |
|---|---|---|
| SDIERQ... | -1.5 | 0.96 |
| SDIEEQ... | -1.4 | 0.85 |
| SIEEAF... | -1.3 | 0.45 |
| SIEAKF... | -1.2 | 0.30 |

| Sequence | LL score | Disc score |
|---|---|---|
| SDIEDF... | -1.4 | 0.88 |
| SDIEAK... | -1.2 | 0.78 |
| STIEAF... | -1.1 | 0.41 |
| STIAKF... | -1.2 | 0.37 |

| Sequence | LL score | Disc score |
|---|---|---|
| SDIERA... | -1.1 | 0.99 |
| SDIERQ... | -1.1 | 0.98 |
| STIEEA... | -1.0 | 0.88 |
| STIEAA... | -0.8 | 0.45 |

**Figure S4:** Generating and selecting proteins for synthesis. (a) A high-level overview of the artificial protein generation process for generic protein family ***f*** using ProGen. ProGen is fine-tuned to a raw sequence dataset of proteins within family ***f***. The resulting model conditionally can generate artificial protein sequences by conditioning on the control tag for family ***f.*** The resulting sequences can be optionally filtered using quality ranking criteria, and top ranked sequences are synthesized and characterized in a laboratory. (b) Overview of pipeline used for lysozyme generation and synthesis. ProGen was fine-tuned to a dataset of known lysozymes across the five target families. Fine-tuned ProGen was then used to generate one million artificial lysozymes conditioning on control tags for the five families. These million sequences were ranked for quality using log-likelihood scores (i.e., model likelihoods) from fine-tuned ProGen, and the probability that an adversarial discriminator predicted that the sequences were real. Since we could only test a limited number of proteins experimentally, we further shortlisted sequences across a range of sequence diversities. For this purpose, the maximum sequence identity to known natural sequences was measured for all artificial sequences. The artificial sequences were then binned into five groups of max ID range, 0.4-0.5, 0.5-0.6, 0.6-0.7, 0.7-0.8 (not shown), and 0.8-0.9 (not shown). The top ranking sequences by discriminator score and log-likelihood score in each bin were selected for synthesis. Sequences were selected one by one in order of rank, and sequences that were more than 80% similar to a previously selected sequence were skipped to ensure diversity.

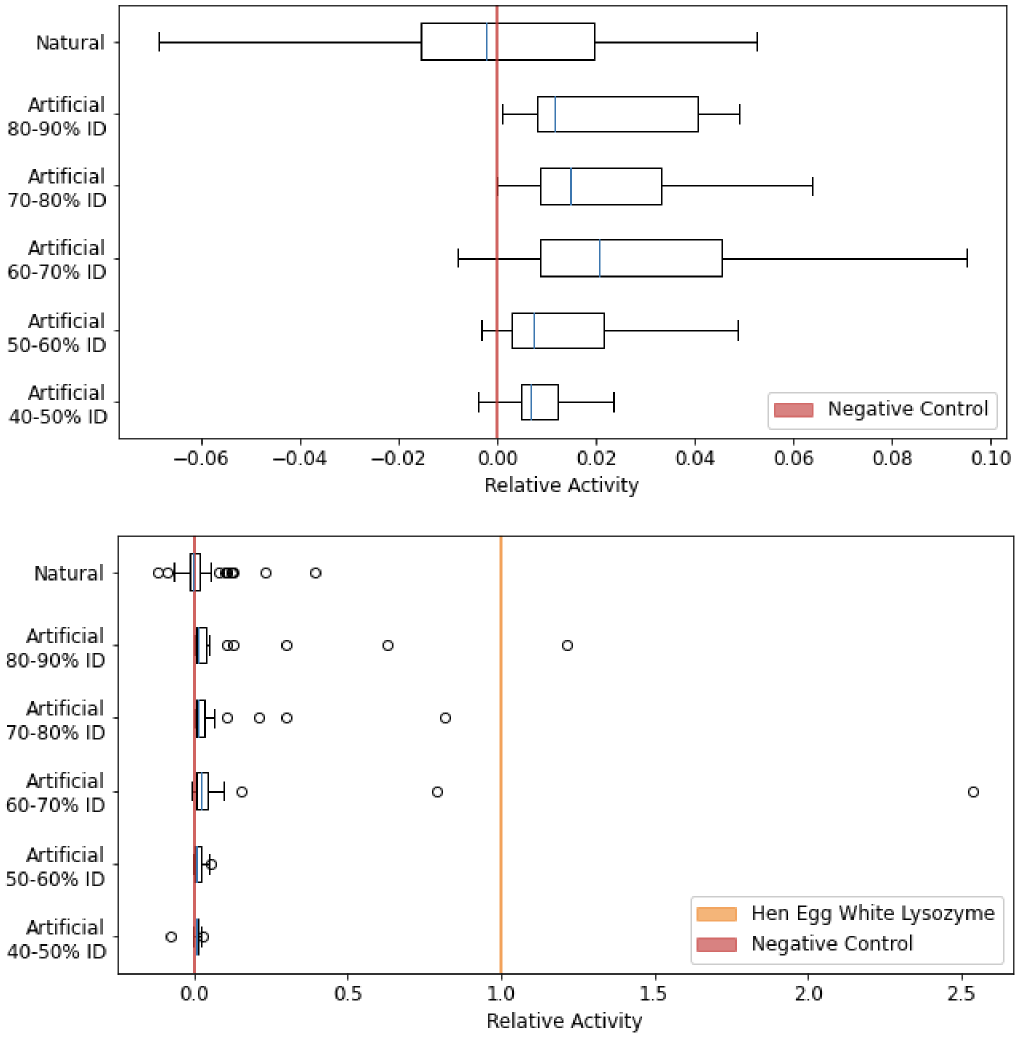

**Figure S5:** Relative activity of natural and artificial lysozymes as measured by the Enzchek kit with the high-throughput *in vitro* translation/transcription (HT-IVT) expression protocol. (Top) The artificial proteins across all max identity bins have a shifted distribution toward more functional values than the natural proteins. The max identity is calculated by finding the closest sequence in any publicly available database of natural proteins. (Bottom) While the majority of natural and artificial proteins are less active than hen egg white lysozyme, there exist outliers out of the one hundred samples, with substantially higher activity.

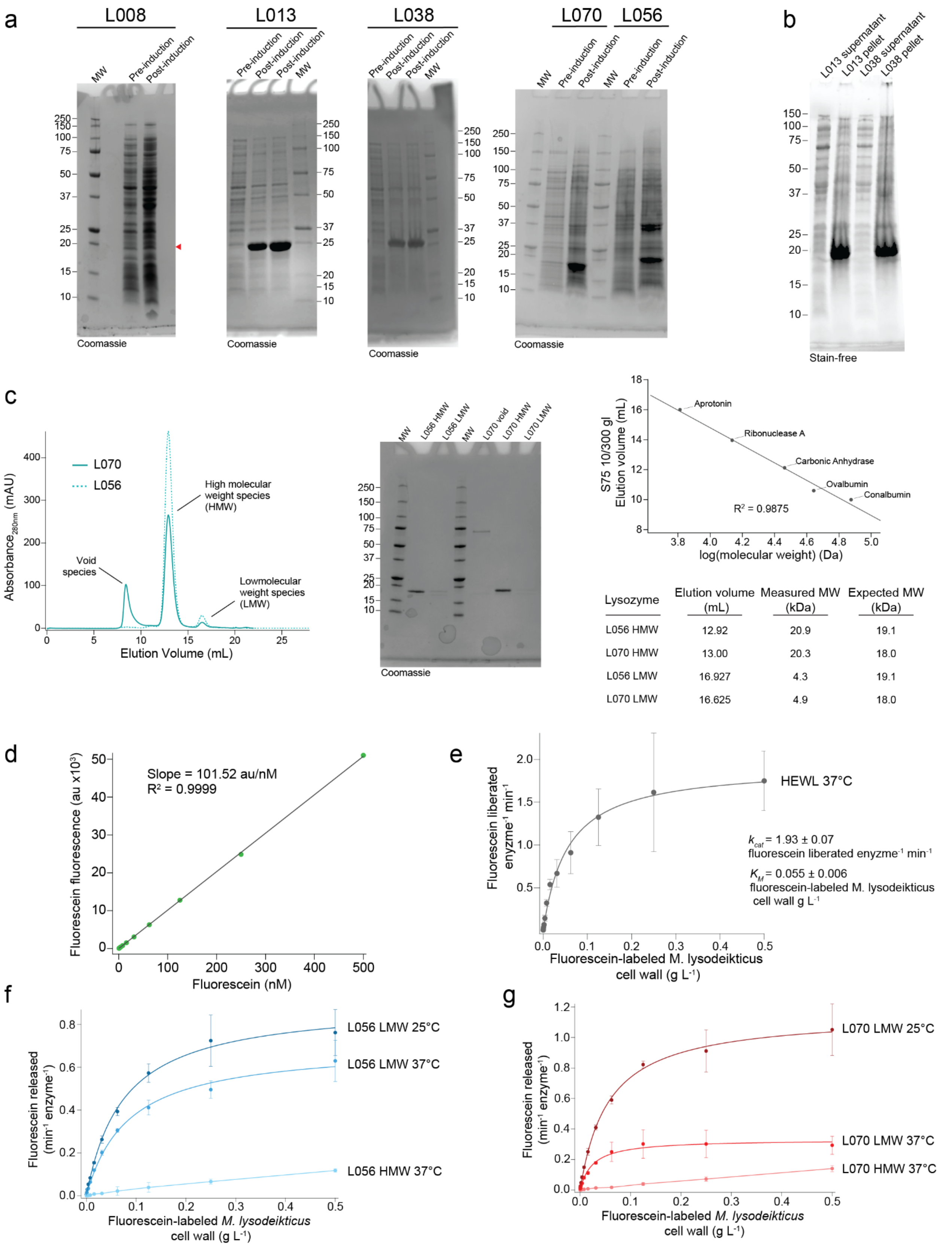

| Lysozyme | Elution volume (mL) | Measured MW (kDa) | Expected MW (kDa) |
|---|---|---|---|
| L056 HMW | 12.92 | 20.9 | 19.1 |
| L070 HMW | 13.00 | 20.3 | 18.0 |
| L056 LMW | 16.927 | 4.3 | 19.1 |
| L070 LMW | 16.625 | 4.9 | 18.0 |

**Figure S6:** (a) Analysis of generated lysozyme overexpression in *E. coli* BL21(DE3) by SDS-PAGE. Red arrow indicates the expected molecular weight (MW) of L008 that was not observed. All other variants tested displayed robust overexpression. (b) Lysate clarification by centrifugation demonstrates that L013 and L038 expressed as insoluble inclusion bodies and were found entirely in the pellet fraction. Stain-free imaging was conducted as per manufacturer protocol (Bio-Rad) on 15-well Any kD mini-PROTEAN gels. (c) Representative size-exclusion chromatography elution profiles of L056 and L070 with relevant peaks labeled (left). SDS-PAGE analysis of different native molecular forms of L056 and L070 indicate that both species are a result of different oligomeric forms and not a cleavage product (middle). Standard curve for S75 10/300 gl (GE; top right) used to estimate molecular weight of the two species of L056 and L070 (bottom right). (d) Fluorescein standard curve utilized in the analysis of initial rates. (e) HEWL Michaelis-Menten kinetics using the fluorescein-labeled *Micrococcus lysodeikticus* cell wall substrate (Molecular Probes EnzChek Lysozyme kit) at 37°C. Points represent the average and error bars the standard deviation of technical replicates ($n$ = 3). Line is resultant of nonlinear curve fitting to the Michaelis Menten model (Eq. 4). (f) Low molecular weight (LMW) L056 Michaelis-Menten kinetics using the fluorescein-labeled *Micrococcus lysodeikticus* cell wall substrate (Molecular Probes EnzChek Lysozyme kit) at 25°C and 37°C. Points represent the average and error bars the standard deviation of technical replicates ($n$ = 3). Lines for LMW L056 data are resultant of nonlinear curve fitting to the Michaelis Menten model (Eq. 4). High molecular weight (HMW) L056 activity against a titration of fluorescein-labeled *Micrococcus lysodeikticus* cell wall substrate (Molecular Probes EnzChek Lysozyme kit) at 37°C could not be fit to Eq. 4 and is represented here as lines connecting points. (g) Low molecular weight (LMW) L056 Michaelis-Menten kinetics using the fluorescein-labeled *Micrococcus lysodeikticus* cell wall substrate (Molecular Probes EnzChek Lysozyme kit) at 25°C and 37°C. Points represent the average and error bars the standard deviation of technical replicates ($n$ = 3). Lines for LMW L056 data are resultant of nonlinear curve fitting to the Michaelis Menten model (Eq. 4). High molecular weight (HMW) L056 activity against a titration of fluorescein-labeled *Micrococcus lysodeikticus* cell wall substrate (Molecular Probes EnzChek Lysozyme kit) at 37°C could not be fit to Eq. 4 and is represented here as lines connecting points.

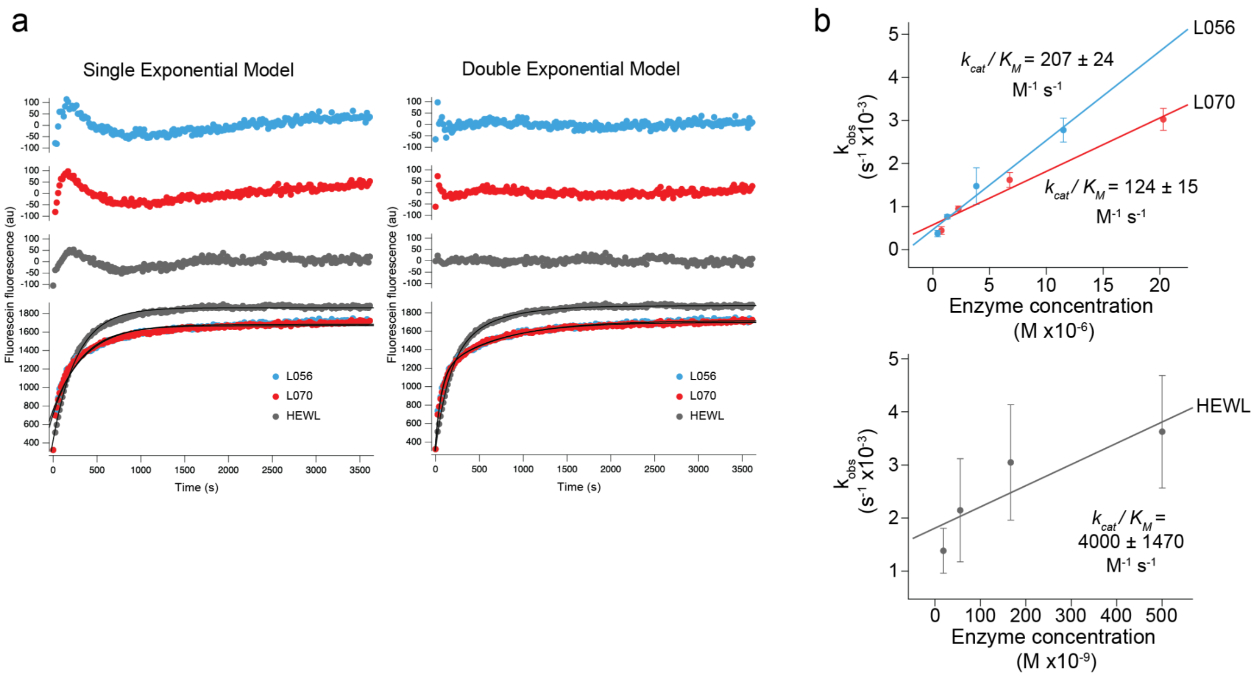

**Figure S7.** Pseudo-first order reaction kinetic analysis. (a) Residual analysis of data fitted by a single exponential model (left) or double exponential model (right; Eq. 6). (b) Linear fitting extrapolation of $k_{cat}/K_M$ from L056 (blue; left), L070 (red; left) and HEWL (grey; right) from Eq. 5. Uncertainty represents standard deviation of fit. Points represent the average and error bars represent the standard deviation of technical replicates ($n$ = 5).

| | $k_{cat}$ / $K_M$ at 37°C ($M^{-1}s^{-1}$) |
|---|---|
| **Hen Egg White Lysozyme** | 4000 +/- 1470 |
| **L56 High molecular weight species** | 207 +/- 24 |
| **L70 High molecular weight species** | 124 +/- 15 |

**Table S4.** Catalytic efficiency of hen egg white lysozyme and monomeric artificial proteins at 37°C. Results indicate L056 and L078 have substantially different Michaelis-Menten constants compared to HEWL and compared to their lower molecular weight counterparts.

| | L056 (7RGR) |
|---|---|
| **Data collection** | |
| Space group | $P2_12_12_1$ |
| Cell dimensions | |
| *a*, *b*, *c* (Å) | 61.15, 68.1, 95.41 |
| α, β, γ (°) | 90, 90, 90 |
| Resolution (Å) | 55.5-2.475 (2.54-2.475) * |
| $R_{meas}$ | 0.1372 (1.75 n/a) |
| $I / \sigma I$ | 13.72 (1.96) |
| Completeness (%) | 99.9 (99.6) |
| Redundancy | 12.8 (12.0) |
| **Refinement** | |
| Resolution (Å) | 2.48 |
| No. reflections | 14740 |
| $R_{work}$ / $R_{free}$ | 0.2525 / 0.2915 |
| No. atoms | |
| Protein | 5393 |
| Ligand/ion | 14 |
| Water | 33 |
| *B*-factors | 62.8 |
| Protein | 69.21 |
| Ligand/ion | 94.2 |
| Water | 55.5 |
| R.m.s. deviations | |
| Bond lengths (Å) | 0.0027 |
| Bond angles (°) | 0.59 |

*Values in parentheses are for the highest-resolution shell.

**Table S5:** Data collection and refinement statistics (molecular replacement) for L056 crystal structure
